# Multi-frame biomechanical and relaxometry analysis during in vivo loading of the human knee by spiral dualMRI and compressed sensing

**DOI:** 10.1101/2023.02.12.528211

**Authors:** Woowon Lee, Emily Y. Miller, Hongtian Zhu, Stephanie E. Schneider, David A. Reiter, Corey P. Neu

## Abstract

**Purpose:** Knee cartilage experiences repetitive loading during physical activities, which is altered during the pathogenesis of diseases like osteoarthritis. Analyzing the biomechanics during motion provides a clear understanding of the dynamics of cartilage deformation, and may establish essential imaging biomarkers of early-stage disease. However, in vivo biomechanical analysis of cartilage during rapid motion is not well established.

**Methods:** We used spiral DENSE MRI on in vivo human tibiofemoral cartilage during cyclic varus loading (0.5 Hz) and employed compressed sensing on the k-space data. The applied compressive load was set for each participant at 0.5× body weight on the medial condyle. Relaxometry methods were measured on the cartilage before (T_1ρ_, T_2_) and after (T_1ρ_) varus load.

**Results:** Displacement and strain maps showed a gradual shift of displacement and strain in time. Compressive strain was observed in the medial condyle cartilage and shear strain was roughly half of the compressive strain. Male participants had more displacement in the loading direction compared to females, and T_1ρ_ values did not change after cyclic varus load. Compressed sensing reduced the scanning time up to 25-40% when comparing the displacement maps and substantially lowered the noise levels.

**Conclusion:** These results demonstrated the ease of which spiral DENSE MRI could be applied to clinical studies due to the shortened imaging time, while quantifying realistic cartilage deformations that occur through daily activities, and that could serve as biomarkers of early osteoarthritis.

## INTRODUCTION

Measuring the mechanical response of knee cartilage by load is important since it is directly related to the load-bearing function of the tissue, and could also serve as an indicator of joint disorders caused by aging and osteoarthritis (OA) [1]–[3]. Cartilage degeneration involves a structural and chemical transition that includes loss of proteoglycan which affects the tissue stiffness [4], [5] thus, the mechanical responses can be a hallmark of tissue degeneration. Enzymatically degraded articular cartilage has shown a decrease in static and dynamic moduli [3]. Also, the shear and elastic modulus measured by indentation has been reported to decrease in degraded cartilage and shows a strong correlation with OA grade and age [1], [2]. Notwithstanding the findings of these studies, the correlation of the acquired biomechanical information with tissue degeneration occurring during the dynamic process the tissue experiences in vivo is rather unclear. This is due to technical challenges including low temporal resolution and the invasive nature of mechanical loading experiments. Joints in the human body experience rapid motion throughout daily activities such as walking and jogging. Thus, analyzing the biomechanics in vivo and under physiologically-relevant loading conditions would greatly help improve understanding of how knee joints degrade over time.

Displacement encoding with stimulated echoes (DENSE) MRI is a quantitative imaging technique that can extract pixel-level displacement fields of the imaged tissue [6]. Given the well-established advantages of MRI for imaging structures in diarthrodial joints (i.e., high spatial resolution, high tissue contrast, full-volume coverage, etc.), DENSE MRI leads to tissue displacement measurements that are potentially accessible in a clinical setting. DENSE MRI has been applied to multiple tissue types, including brain and heart. Pixelwise regional and whole brain tissue motion have been measured using DENSE MRI potentially being indicative of brain defects [7], [8]. Intervertebral disc and cartilage, which routinely experience mechanical loading, have also been analyzed by DENSE MRI to quantify displacement and strain during compression [9], [10].

To enhance the temporal resolution of DENSE MRI, non-cartesian sampling approaches have been employed. Spiral image acquisition samples data in a spiral interleave trajectory on k-space using sinusoidal gradient waveforms [11]–[13]. Unlike cartesian scanning [14], [15], spiral image acquisition has advantageous for dynamic imaging because the center of k-space, which carries the most information, is sampled with every interleave and only a few numbers of interleaves are needed to cover a large amount of k-space data. Previous work has shown that the signal intensity of spiral scanning increases compared to cartesian scanning due to the shorter TE and readout efficiency [16], [17]. Spiral DENSE MRI has been successfully used on cardiac tissue to analyze the displacement and strain in the left ventricle during the cardiac cycle [18], [19]. These studies clearly leverage the high temporal resolution of spiral acquisition since the data and analysis were conducted on the rapid cardiac cycle. Recently, we have demonstrated spiral DENSE MRI on bovine joints with a temporal resolution of 40 ms during compression and have found differences between intact and defected bovine joints [20]. Also, we have successfully measured the viscoelastic response of cartilage using spiral DENSE MRI.

Compressed sensing (CS) is a technique which can reconstruct the image with significantly under-sampled data [21], [22]. This is due to three basic principles: sparsity of the data, random data collection, and iterative non-linear reconstruction that implements the sparsity and consistency with the collected data. Applying CS to MR images is feasible since MRI captures k-space where the data is sparse. Also, spiral scanning generates noise-like artifacts in the imaging domain which is analogous with effects caused by random sampling [22], and can be reduced by non-linear reconstruction methods. CS has been used in combination with spiral acquisition for imaging cardiac and liver tissue [23]–[25], demonstrating accelerated scanning speed permitting breath-hold imaging and showing comparable image quality with other advanced methods such as SENSE [26]. DENSE MRI generally collects data over multiple cycles of tissue motion for ample signal intensity which leads to lengthy imaging time. Therefore, DENSE MRI can greatly benefit from CS by reducing the scanning time which is more tolerable for participants and induces less motion artifacts for maintaining a static posture in the reduced scanning time [27].

In this paper, we acquired high frame rate (25 frames/s) displacement and strain maps on healthy human in vivo cartilage using spiral DENSE MRI during varus load. The loading frequency was 0.5 Hz, comparable to walking cadence. A gradual increase of displacement and strain was observed in the medial condyle which is consistent with visual observation and coherent with the intent of the loading device. In addition, we applied CS on the under-sampled (low average) k-space data and obtained comparable image quality with fully-sampled (high average) data saving up to 25-40% of imaging time. Quantitative MRI measurements (T_1ρ_, T_2_) showed similar values with previous studies and no change (T_1ρ_) after varus loading was observed which entailed minimum discernable effects from the varus load. We believe our approach using spiral DENSE MRI combined with CS can easily be used in a clinical setting which can provide useful biomarkers predicting early degenerations of tissue.

## METHODS

The knee imaging protocol consisted of a series of sequences (Figure 1) including multi-slice double echo steady state (DESS), quantitative T_1ρ_ and T_2_ measurements, followed by DENSE MRI during varus loading on the knee joint. An additional quantitative T_1ρ_ measurement was acquired after the loading protocol. All imaging was performed using a clinical MRI system (3 T; Siemens Prisma^fit^) with a 15-channel knee coil (Tx/Rx Knee 15 Flare Coil, QED, LLC). DENSE k-space data was extracted and analyzed off-line for CS. All experiments were approved by the Institutional Review Board at the University of Colorado, Boulder.

**Figure 1.**
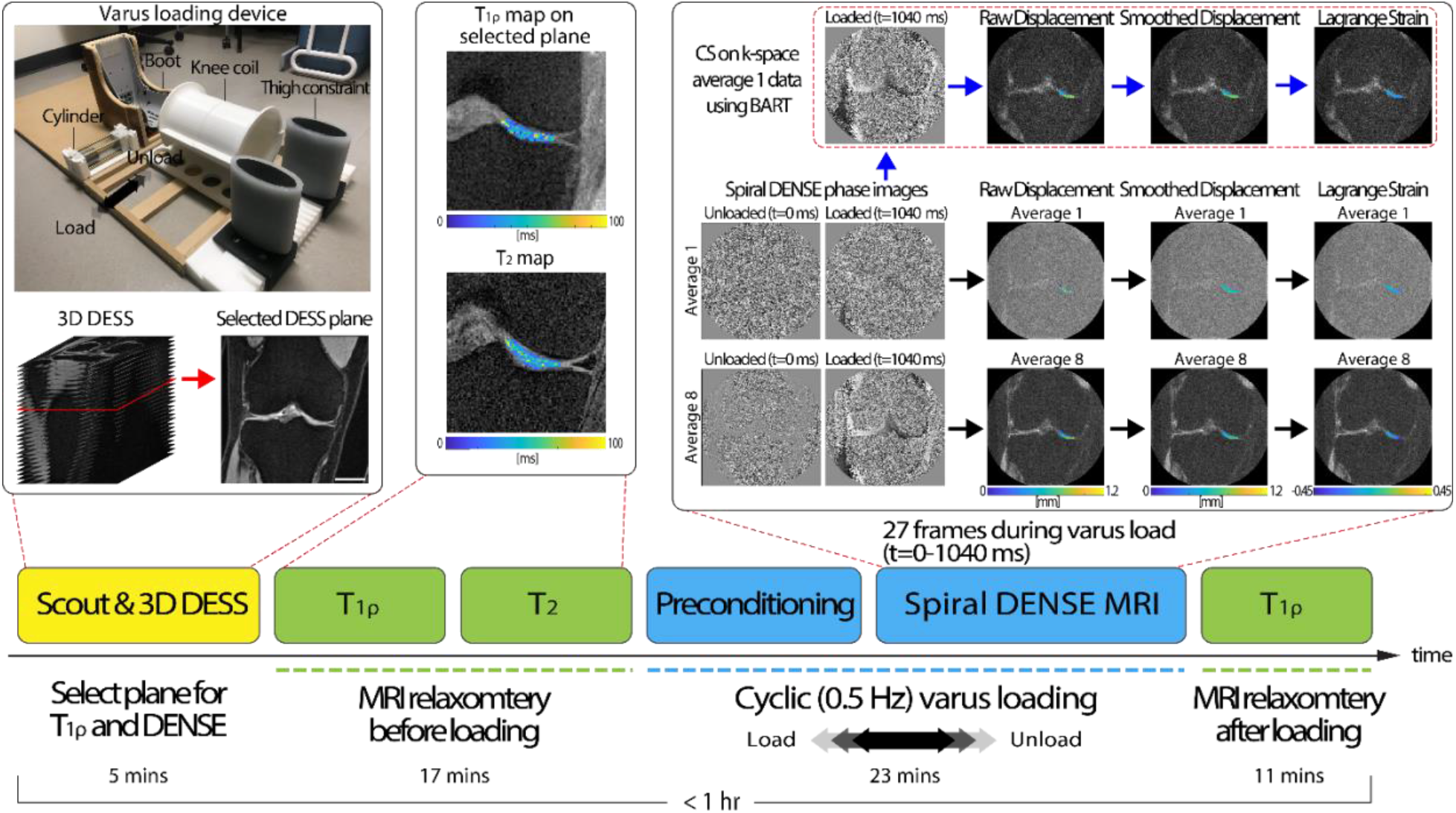
The experimental design for acquiring relaxometry measures (T_1ρ_, T_2_) and spiral DENSE MR images during rest and varus loading. We implemented a pneumatic loading device which consists of thigh constraint, cylinder, and a boot section. Participants were positioned in a supine position inside the MRI bore while the knee coil collected MR images. The thigh constraint held the thigh while the pneumatic load from the cylinder pulled the boot to apply varus load on the knee. Varus load was applied in a cyclic fashion consisting of 1 s of load followed by 1 s of unload. On a static rest condition with no varus load, the experiment started with obtaining fast gradient echo MR images (scout) for localization then subsequentially 3D DESS acquisition to select a plane where there was a large tibiofemoral contact area in the medial condyle. After DESS acquisition, relaxometry measurements (T_1ρ_, T_2_) were ran. Next, cyclic varus load was applied and after 8 minutes of preconditioning, spiral DENSE MR images were collected. The saved spiral DENSE MR phase images in *x* and *y* were converted to displacement fields in the selected ROI and smoothed prior to calculating Lagrange strain. To apply CS, k-space data was saved at the scanner and was inputted to BART which reconstructed CS phase images. On the reconstructed CS image, the equivalent process to calculate strain was applied. The displacement and strain calculation were conducted off-line. Lastly, T_1ρ_ measures were collected after the varus loading was stopped. The images shown were collected on a 29-year-old male subject. Scale bar=25 mm.

### Varus Loading on Human Knees

Eight subjects (4 males/4 females, age=20-29, BMI=22.86±1.92) with no history of cartilage damage were recruited. Varus loading was applied to the knee joint through a lateral force at the participant’s ankle resulting in a compressive load on the medial tibiofemoral cartilage. Loading was applied to the ankle using a pneumatic actuator calibrated to produce the equivalent to 0.5× the participant’s body weight (BW) [10], [28] (Figure 1). The loading device was designed to fully constrain lateral motion of the thigh while applying lateral loading using a pneumatic cylinder to brace affixed to the ankle. Participants were positioned supine with neutral knee flexion. The load at the knee was estimated using a moment balance, the controlled force applied at the ankle, and moment arms extending from the knee joint (centerline) to the load application point at the ankle (measured by tape measure), and additionally from the knee joint center to the center of the condyle (measured on DESS images). Mounted foam was utilized to minimize translation motion of the knee inside the coil. Among the DESS slices, a plane with large tibiofemoral contact area in the medial condyle was selected (Figure 1). For the experiment, varus loading was applied in a cyclic fashion (0.5 Hz) consisting of 1 s of loading followed by 1 s of unloading, while there was a 100 ms delay prior to each loading cycle. To further analyze the performance of CS on different image averages, we recruited additional seven subjects (3 males/4 females, age=26-35, BMI=21.39±3.23) with healthy knees and applied the equivalent cyclic varus load for imaging.

### Multi-frame Spiral DENSE MRI

Prior to DENSE MRI acquisition, we applied cyclic varus load without imaging (8 mins) to gain a quasi-steady state response of cartilage [29]. Subsequentially, we collected multi-frame (27) coronal plane images using spiral DENSE MRI [18]–[20] with the following parameters: temporal resolution=40 ms, interleaves=10; TE/TR=2.5/20 ms, spatial resolution=360×360 µm^2^, slice thickness=1.7 mm, and in-plane displacement encoding gradient=0.64 cycles/mm. All images were collected while the cyclic varus load was applied on the knee joint and in a single slice matching with the preselected DESS slice. Two separate data sets were collected (image average 1, 8) to control the signal-to-noise ratio (SNR) and imaging time. Imaging time increased linearly with averages, with 1 and 8 averages requiring approximately 1.7 and 13.6 mins, respectively. The average 1 k-space data was saved for the CS analysis. For the second group of participants (n=7), we conducted one spiral DENSE MRI with 8 averages and saved the k-space data.

### Displacement and Strain Calculation

Tibiofemoral contact areas were manually segmented using ImageJ (1.53t) and were utilized as binary masks. A fixed region of interest (ROI) on the medial condyle was drawn on frame 1 (t=0 ms) and manually translated along the movement of the joint on each frame. Phase images were used to compute *x* and *y* displacement maps within the ROIs [20] and subsequently smoothed by a locally weighted scatterplot smoothing filter for 30 cycles with a span of 30 [30]. Next, the displacements were smoothed with a Gaussian filter in time with a kernel size of 3 and SD of 2 [20]. Finally, the smoothed displacement maps were used to calculate in-plane Green-Lagrange strains (Figure 1).

### Relaxometry (T_1ρ_, T_2_)

T_1ρ_ and T_2_ measures were obtained on the knee joint of all participants (n=8) while in static rest condition prior to cyclic varus load (Figure 1). T_1ρ_ measurements were made using a magnetization-prepared gradient echo sequence [31] with the following acquisition parameters: TE/TR=3/6 ms, spin-lock frequency=500 Hz, spin-lock durations=50, 100, 150, 200 ms, FOV=90×90 mm^2^, spatial resolution=700×700 µm^2^, slice thickness=3.5 mm, slice number=1 (on the selected DESS slice), flip angle=10°, image average=2. T_1ρ_ relaxation maps were calcualted by fitting the four MR intensity images to a monoexponential relaxation model using non-linear least squares. Voxel fits with coefficient of determination (R^2^) less than 0.66 were removed [32], [33]. T_2_ mesurements were made using a multi-echo spin echo sequence with the following acquisition parameters: TEs=13, 26, 39, 52, 65, 78 ms, TR=1290 ms, FOV=80×80 mm^2^, spatial resolution=420×420 µm^2^, slice thickness=1.7 mm, slice number=13, flip angle=180°, image average=2. T_2_ relaxation maps were calculated by fitting multi-echo signal intensities to a monoexponential using non-negative least squares [34]. The tibiofemoral contact area on the medial condyle was selected for the ROI and the spatial average (SA) of the pixels in the ROI was calculated. Both measurements used fat suppresion. T_1ρ_ measurements were repeated after the cyclic varus loading to see the affects of varus load on the T_1ρ_ values.

### Applying CS on Spiral DENSE MRI

The k-space binary files were input to a customized MATLAB (Mathworks, R2019b) code to extract the k-space values along the spiral trajectories for each channel, image average, and frame. We then applied CS analysis using the open-source toolbox, Berkeley Advanced Reconstruction Toolbox (BART), parallel imaging and CS command to reconstruct the images [35]. The inputs on BART were the spiral trajectory, channel sensitivity map and k-space data along the interleaves and the output was the reconstructed magnitude and phase images. The coil sensitivity map and image reconstruction were generated by using the ESPIRiT method in BART. We used l2 regularization (λ=0.01) and all processes underwent five iterations [27]. This process was applied to each individual average (1-8) and frame (1-27). Subsequently, image averaging of the CS reconstructed images was performed on both the magnitude and phase images. All images were then used for displacement and strain measurements (Figure 1).

### Statistical Analysis

The SNR was measured by first selecting the cartilage in the medial condyle and background region in the DENSE MR magnitude image. Subsequently, we took the mean signal in the cartilage and then subtracted by background noise and later divided by the SD of the signal in the background [36]–[38]. All datasets were analyzed using a linear mixed effects model (nlme package, Version 3.1-140) with type II or III sum of squares ANOVA (car package, version 3.0-12) in R (RStudio, Version 1.2.1335; R, Version 3.6.1). In all models, participants were considered a random effect. Normality assumptions required for ANOVA were validated by testing the residuals of the model with the Shapiro-Wilk test and visual examination of qq-plot and residual plots. Post-hoc tests were performed with the emmeans package using Tukey’s HSD corrections for multiple comparisons and interaction terms (Version 1.4.3.01). For SA displacement and strain datasets, the raw data showed a binomial distribution which allowed the data to be classified into two groups of loading status: loading (t=80-440 ms) and loaded (t=480-1040 ms). The model tested the relationship between loading status and gender. For the T_1p_ and T_2_ dataset, the model tested the relationship between gender and before and after loading. For the non-CS (NCS) and CS displacement and strain datasets, the models tested the relationship between averages and processing (NCS or CS). Pearson correlation coefficient was calculated in R using the built-in stats package. Assumptions of normality and linearity were validated prior to testing the relationship.

## RESULTS

### Displacement and Strain Analysis on Cartilage from Spiral DENSE MRI

The displacement maps captured on the medial condyle from spiral DENSE MRI showed a gradual change in *x* and *y* with time during the applied varus load (Figure 2A). The knee experienced translation within the knee coil due to the varus load, specifically towards the medial condyle and femur direction. This trend was reflected in the displacement fields (Figure 2C; ROI labeled in Figure 2B) where the four representative time points in Figure 2C showed a moderate increase in displacement with time. More displacement was observed in *x* reaching almost 5 mm compared to *y* which was less than 0.6 mm on a representative subject (Figure 2). There were also internal variations within the displacement maps which resulted in strain values (Figure 2C). E_*yy*_ had the maximum amount of strain magnitude which reached negative 0.035 on the representative subject indicating compression while the maximum strain magnitudes in E_*xx*_ and E_*xy*_ were 0.0165 and negative 0.0253, respectively. Compressive strain was induced by the high displacement *y* values in the lower right corner and low values in the upper left corner within the ROI which was caused by the varus loading. Strain values also showed a steady change while the varus load was being applied. All participants (n=8) showed a similar pattern in the displacement and strain fields with little differences in magnitude.

**Figure 2.**
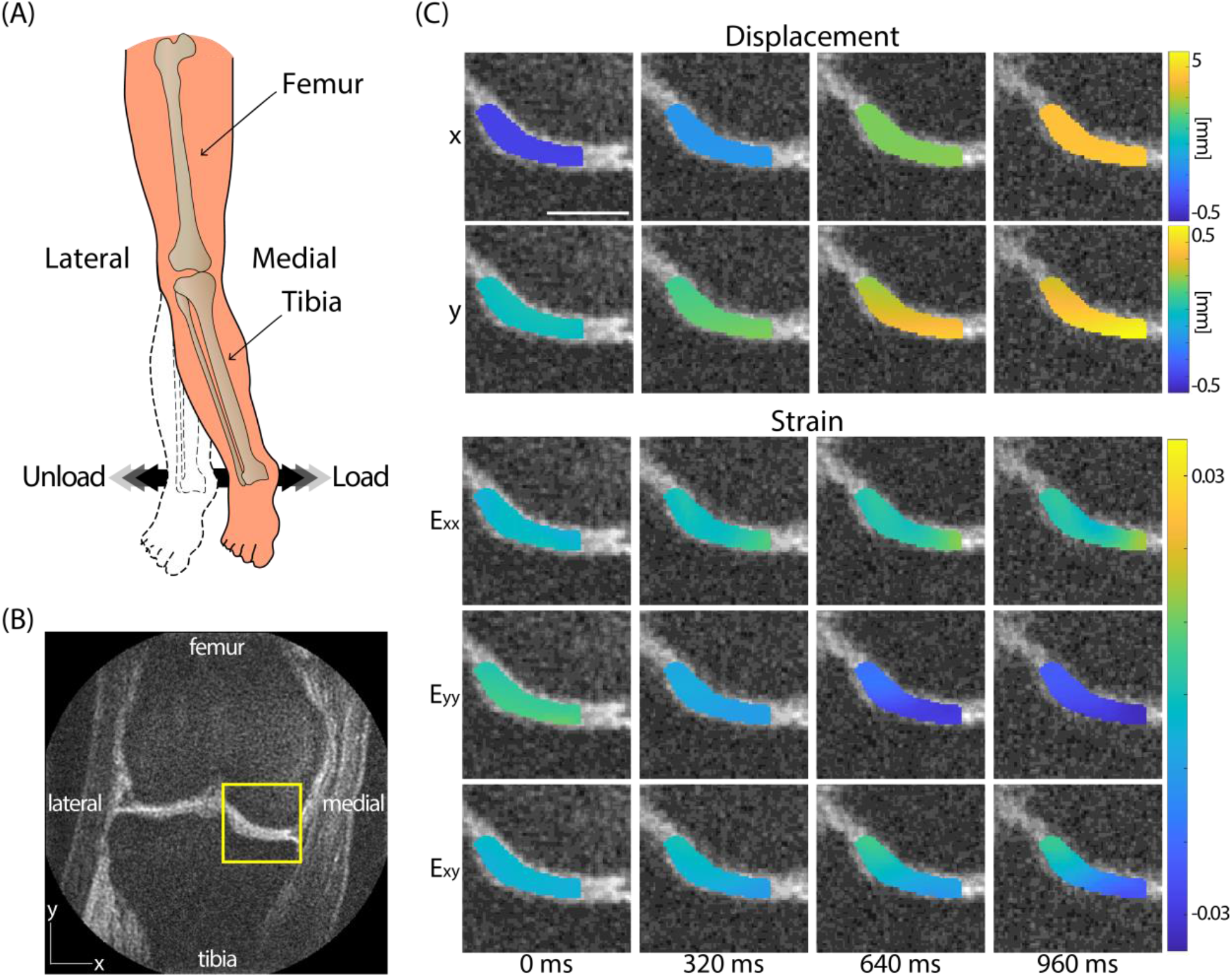
Spiral DENSE MRI during varus load showed time-course (40ms; 25 frames/s) displacement and strain maps in the articular cartilage on the medial condyle. (A) The varus load applied on ankle was controlled to be equivalent to 0.5× BW of compressive load on (B) the medial condyle shown in the representative image. (C) The displacement in both *x* and *y* showed a gradual increase where more displacement was observed in *x* due to the translation occurring in the knee coil. Strain maps also displayed a gradual change in strain. E_*yy*_ showed a pronounced compression occurring while mild tension in E_*xx*_ and shear strain, E_*xy*_. The images were obtained on a 26-year-old female subject. Scale bar=15 mm.

The SA displacement and strain values within the ROI averaged for all participants (n=8) quantitatively showed a gradual transition over time (Figure 3A). The SA displacements were close to 0 mm at t=0 ms and sharply increased starting from around t=160 ms through 760 ms and then moderately increased until 1040 ms. The gradual increase in displacement at early time points was in part due to the load in the actuator having a short period to ramp up before reaching the desired load [20]. The maximum SA displacements were close to 5 and 0.5 mm for *x* and *y*, respectively. Analogous to displacement maps, SA strain values also showed a sharp increase near t=160 ms. As expected, the SA E_*yy*_ showed a compressive strain (maximum -0.0925) indicating that the varus load was effectively being applied to the knee. The SA E_*xx*_ and E_*xy*_ were in the positive range where the maximum magnitudes were less than half of SA E_*yy*_. Positive E_*xx*_ indicates tension which was reasonable due to the Poison effect. SA E_*xy*_ showed a slight decrease in strain starting from t=640 ms (0.0274±0.0329) until 1040 ms (0.0176±0.0294). In SA displacement *x, y*, and E_*yy*_, there were significant differences between loading status (loading, loaded) (*P*<0.0001 for displacement *P*<0.05 for E_*yy*_) (Figure 3A). A significant difference (*P*<0.0001) was detected between gender in displacement *x* (Figure 3B). In the female group, all SA strains reached a peak value at t=560 ms and either showed a steady decrease or no increase in magnitude whereas in the male group there were a continuous increase. The decrease of strain in the female group was mostly pronounced in SA E_*xy*_.

**Figure 3.**
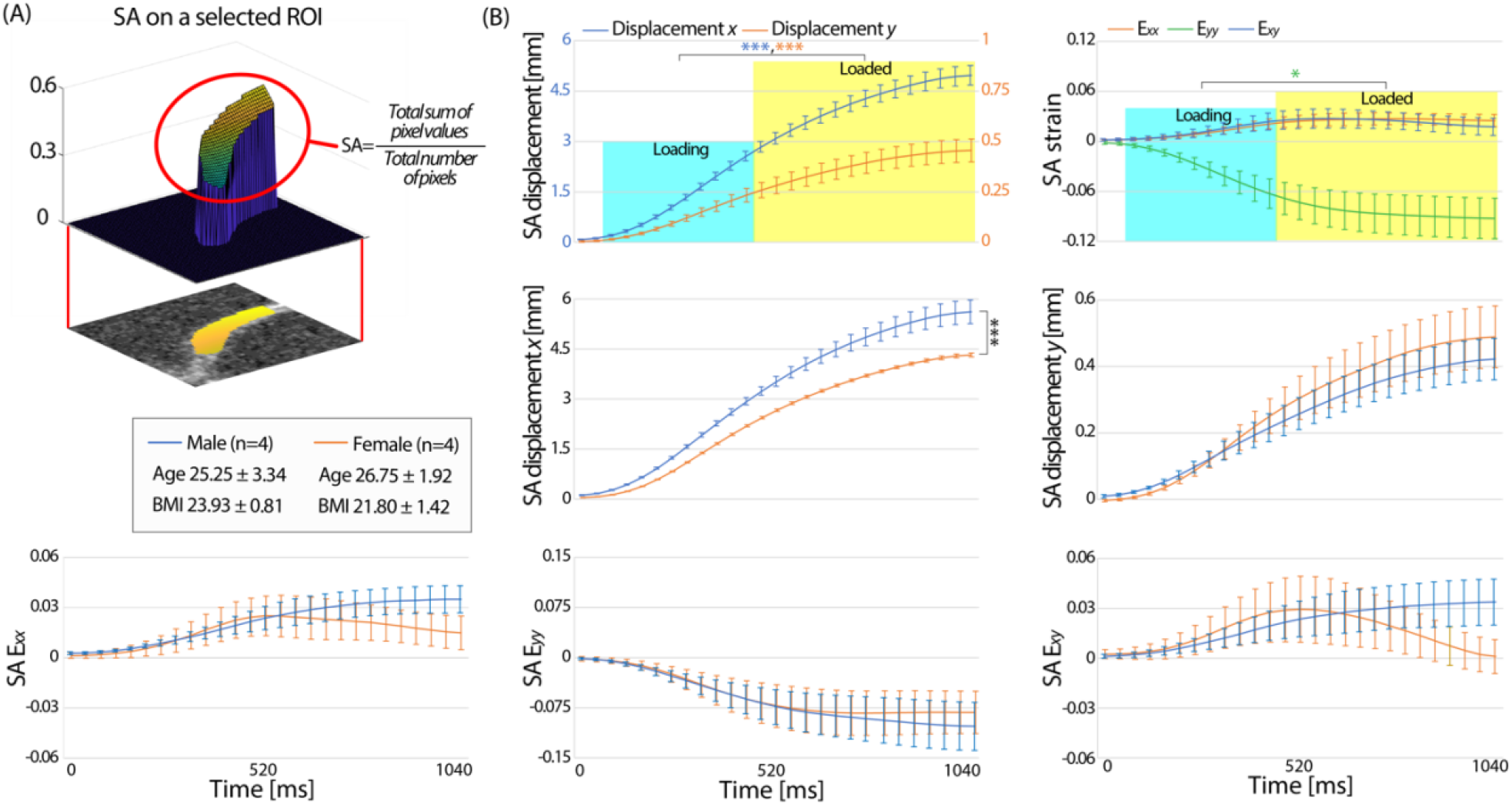
The SA displacement and strain within the ROI showed a gradual transition with time on all participants from the initially recruited group (n=8). (A) All subjects from the former group, showed a gradual increase of SA displacement and strain with time where SA displacement *y* was more than an order of magnitude higher than SA displacement *x*. Compressive strain in SA E_*yy*_ and tension in SA E_*xx*_ was observed on all subjects which validated the cartilage being compressed due do the varus load and spreading in the perpendicular direction to the loading direction. There was a significant difference in the loading status (loading, loaded) for displacement *x, y*, and E_*yy*_. (B) Male subjects had significantly more SA displacement in *x* compared to females. There was no significant difference between gender in SA displacement *y*. The SA strain data in the male group showed a steady increase with time while the strain on females did not increase or decreased after t=560 ms in both normal and shear strain. This resulted in no significant difference in loading status for E_*xx*_ and E_*xy*_. Error bars are the standard error of the mean. \**P*<0.05, \*\*\**P*<0.0001.

### Relaxometry Measurements (T_1ρ_, T_2_)

The measured T_1ρ_ and T_2_ in this study was comparable with previously published studies on subjects with healthy knees [39]–[41] and we found no significant difference between gender or before and after varus load. The T_1ρ_ measured on the cartilage before cyclic loading was 50.47±1.23 ms and 45.43±2.54 ms for the male and female group, respectively (Figure 4A). There was no significant difference found after varus loading in T_1ρ_ (*P*>0.05). Male participants had mildly higher T_2_ values (38.24±0.58 ms) compared to female participants (34.33±4.07 ms) but was not significantly different (Figure 4B). There was no correlation found between T_1ρ_, T_2_ and the strain measured by spiral DENSE MRI (*P*>0.05).

**Figure 4.**
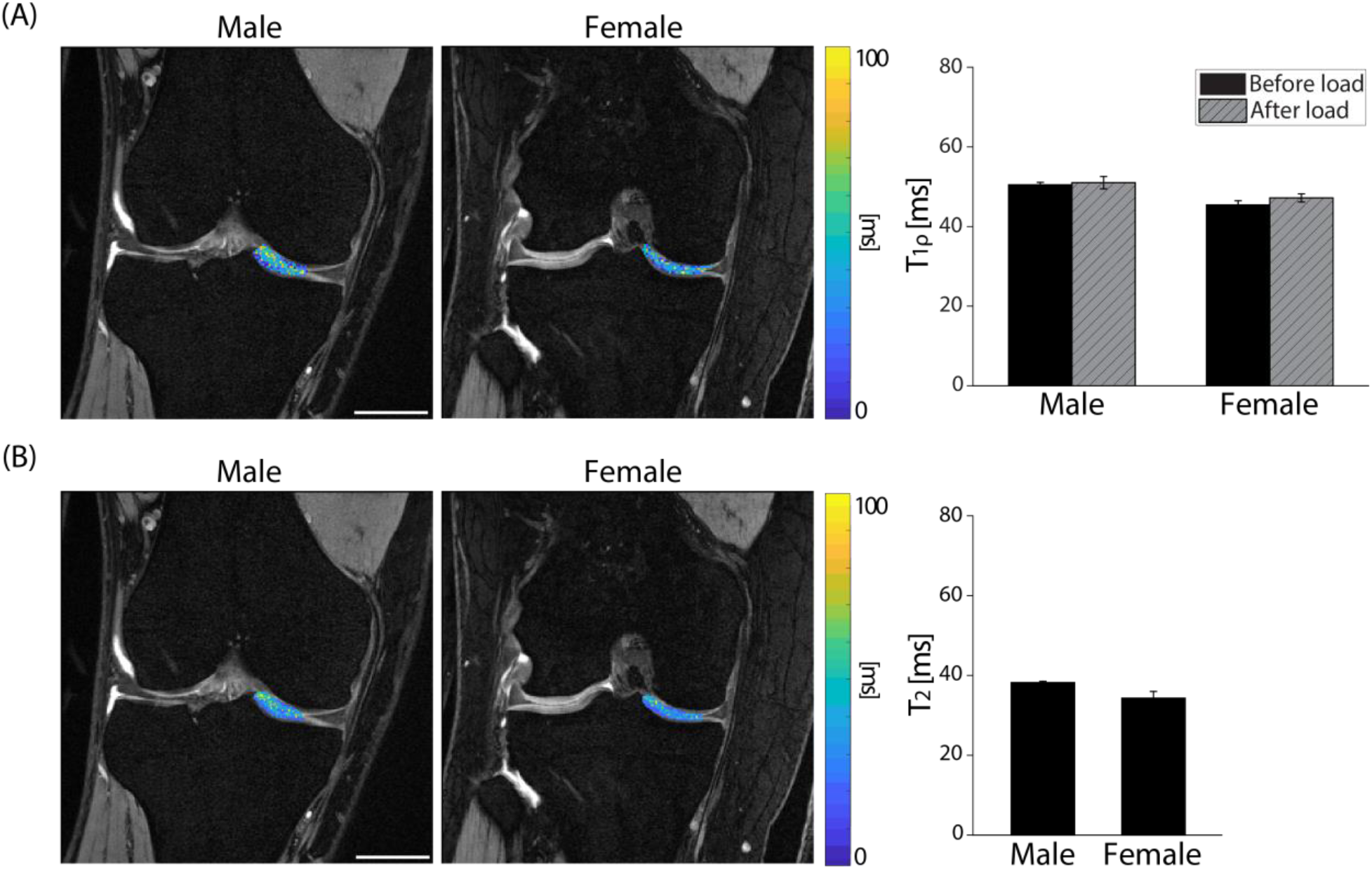
T_1ρ_ and T_2_ relaxometry measurements on the knee cartilage generated pixel-scale maps while no changes were observed after varus loading (T_1ρ_) or between gender (T_1ρ_, T_2_). (A) T_1ρ_ pixel values were overlayed on DESS images collected on a representative male (27-year-old) and female (29-year-old) subject. After cyclic loading, the average T_1ρ_ values slightly increased on both male and female groups but were not statistically significant (*P*>0.05, effect size=0.373). Male and female T_1ρ_ values were in a similar range (*P*>0.05, effect size=0.407). (B) T_2_ values showed an analogous trend with T_1ρ_ where there was no significant difference in gender (*P*>0.05, effect size=0.146). Error bars are the standard error of the mean (n=4). Scale bar=25 mm.

### Analysis on the Effects of CS

The effect of applying CS on average 1 data was distinct, substantially improving the displacement and strain maps to be similar with the average 8 data (Figure 5A,B). Average 1 DENSE MR images resulted in noisy displacement maps showing spiked values in some pixels necessitating the need of smoothing. Even after smoothing, the displacement maps showed a discrepancy compared to the average 8 data. After applying CS, the displacements not only smoothened but also became comparable with the average 8 data. A similar trend was observed in the strain maps, specifically where the compressive strain in E_*yy*_ at t=1040 ms was not pronounced in the average 1 data in contrast with the average 8 and average 1 - CS data showing larger compressive strain. The SNR was significantly larger (*P*<0.0001) in the average 8 data and also in the average 1 - CS data (Figure 5C).

**Figure 5.**
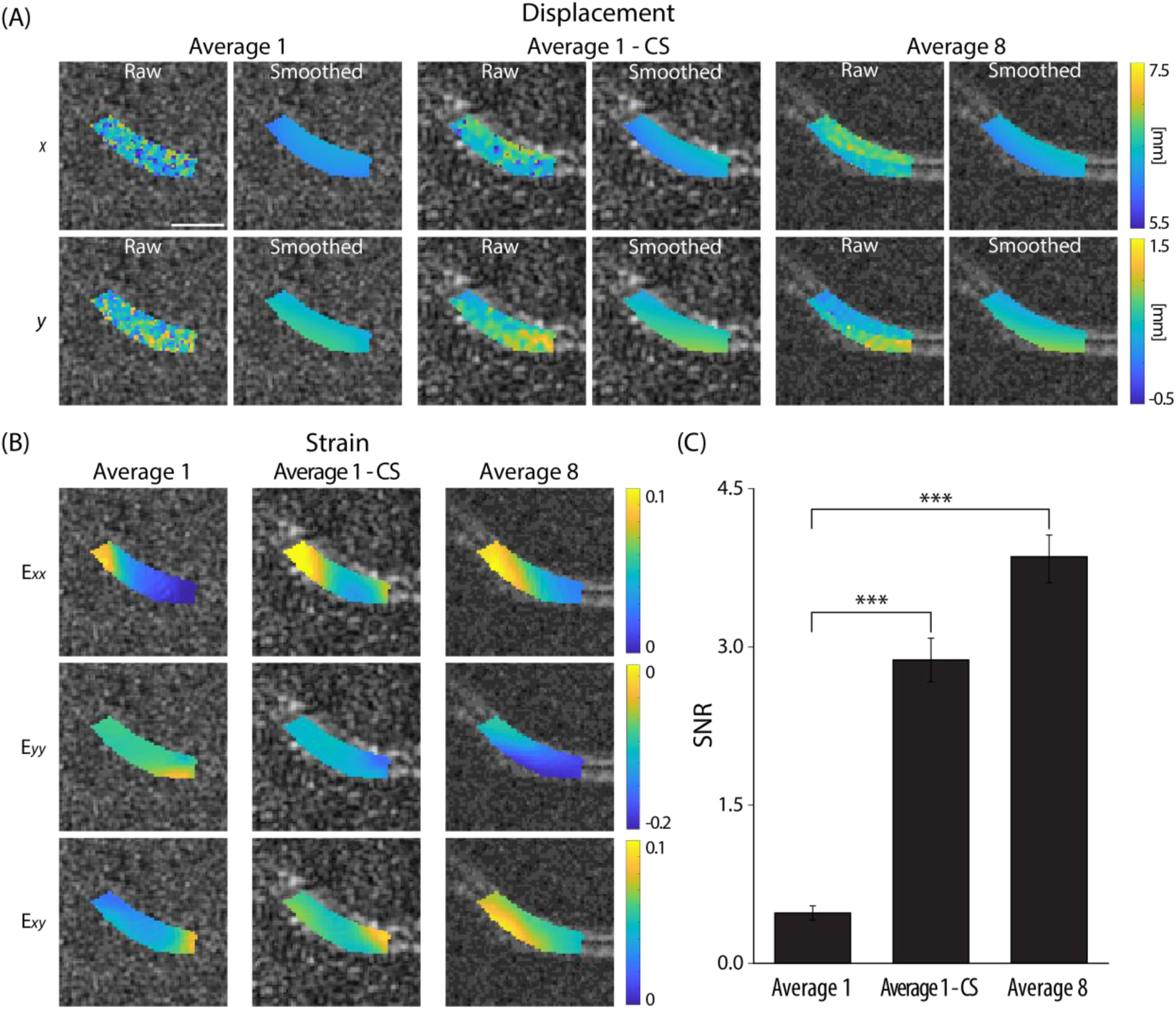
Applying CS on the average 1 k-space data enhanced the accuracy of the biomechanical results at a varus loaded time point (t=1040 ms, frame 27). (A) Displacement maps were compared on image average 1 and average 8 where the former showed higher noise levels resulting in discrepancy between the two averages strongly on the raw displacements prior to smoothing. Applying CS on average 1 data (average 1 - CS) did reduce the noise and error. (B) Strain maps showed a similar trend as the displacement maps where compressive strain at E_*yy*_ became more pronounced after applying CS, and the values were more comparable to the average 8 data. (C) The SNR measurements showed a significant increase in average 1 - CS compared to average 1. All images shown were collected on a 25-year-old male subject. Scale bar=10 mm. \*\*\**P*<0.0001.

The power of CS improving the image quality was persistent on all time-course images (Figure 6). To visualize the difference between the three reconstruction methods (average 1, average 1 - CS, and average 8), SA raw displacement and strain data versus time were plotted as shown in Figure 6A,B. The difference was more substantial in average 1 compared to average 1 - CS. We calculated the average absolute difference in SA displacement and strain of average 1 and average 1 - CS with respect to average 8 across all 27 frames and repeated on all subjects (n=8) and defined them as the SA displacement and strain error (Figure 6, bar graph indicating mean±standard error, n=8). Both SA displacement and strain error improved in average 1 - CS showing a significant difference (*P*<0.05) compared to average 1.

**Figure 6.**
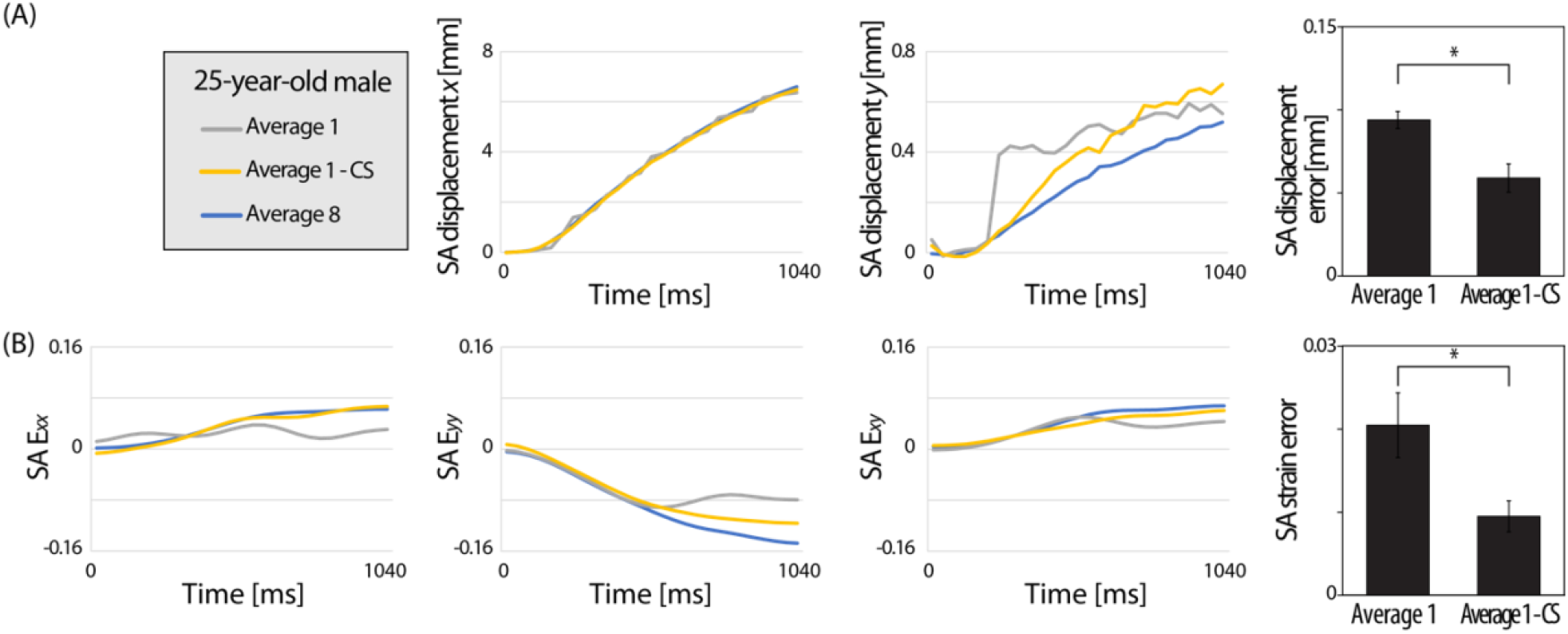
CS improved the correspondence of the SA raw displacement and strain values between the average 1 and average 8 data on all time points. (A) SA displacement and (B) SA strain versus time plots captured on a representative subject (25-year-old, male) visualized the average 1 - CS data approaching the average 8 data points. The error between the reconstruction methods was quantified as the absolute average difference for each time point and averaged for all eight subjects. Significantly reduced SA displacement and strain errors were observed after applying CS. Error bars are the standard error of the mean (n=8). \**P*<0.05.

The impact of CS was explored on images acquired on different averages where there was a consistent improvement on signal intensity and accuracy of displacement and strain values after applying CS on low average data while the improvement due to image averaging was not as strong. The performance of CS was analyzed on the later recruited group (n=7) on different image averages. As expected, more image averaging resulted in less noise, higher signal, and smoother raw displacement maps (Figure 7A). The raw displacement maps on the CS data showed a smoother pattern and higher signal starting from average 1 and moderately increased with more averages. To quantify the effects of CS on each image average, we calculated the SNR and root mean square error (RMSE) of the displacement (*x, y* averaged) and strain (E_*xx*_, E_*yy*_, E_*xy*_ averaged) compared to the average 8 data to account for the difference in each pixel (Figure 7B,C). SNR consistently enhanced by averaging for both NCS and CS groups where the CS average 2 exceeded the SNR of NCS average 7. For raw displacement, the RMSE reduced with more averaging where the two groups met between average 5 and 6. When calculating the RMSE with respect to the smoothed displacement of average 8, the CS group had a constantly lower RMSE in all averages indicating the smoothing effect of CS. The RMSE for raw strain (calculated from raw displacement) showed that the CS group had lower RMSE in all averages when comparing for both raw and smoothed strain (from smoothed displacement) of average 8. Interestingly, for the RMSE of smoothed strain, the NCS and CS group met between average 3 and 4.

**Figure 7.**
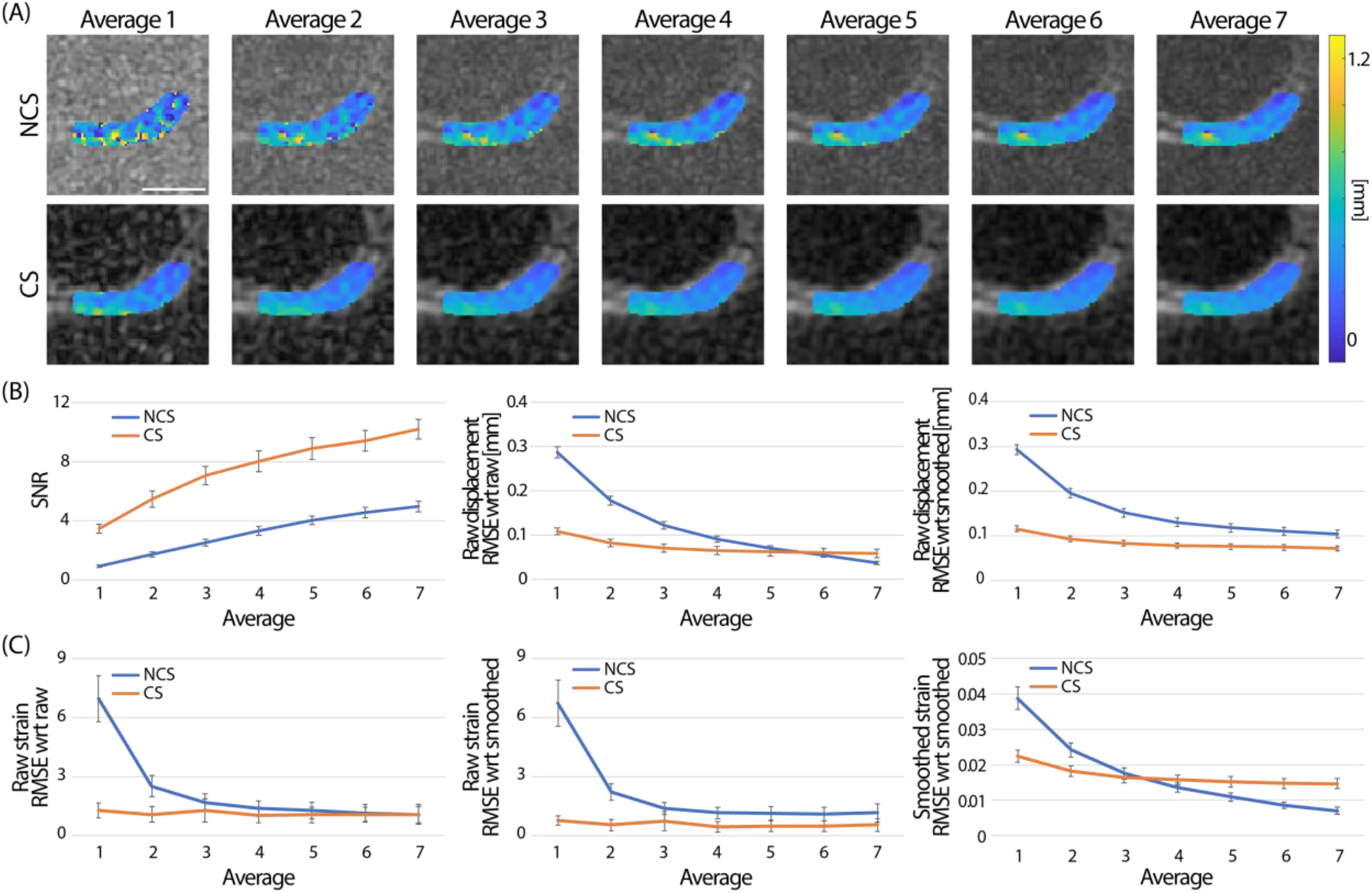
Image averaging and CS improved the signal intensity and smoothness on displacement and strain maps. (A) Raw displacement *y* maps (t=1040 ms) were compared qualitatively between images reconstructed on different averages. With more averaging the smoothness increased and noise level decreased. Images shown were collected on a 25-year-old female subject. Applying CS did show a similar effect where the displacement maps had less pixels with errors and were steadily smoothened with more averaging. (B) Quantitatively, the SNR did increase in both non-CS (NCS) and CS groups with more averaging and CS average 2 group surpassed the SNR of NCS average 7. RMSE of the displacement gradually decreased in the NCS group. RMSE at the CS group substantially improved compared to the NCS at lower image averages and moderately improved with more averaging. The two groups met between average 5 and 6. When comparing the raw displacement with respect to (wrt) the smoothed displacement of the average 8 data, CS showed significantly lower RMSE compared to the NCS group in all averages. (C) A similar trend was observed in strain. When using the raw strain (calculated from raw displacement), the RMSE was lower in the CS group for all averages. This result was consistent for either comparing to raw strain or smoothed strain (calculated from the smoothed displacement) of the average 8 data. When calculating the RMSE of the smoothed strain, NCS and CS met between average 3 and 4. Displacement *x, y*, and all strains (E_*xx*_, E_*yy*_, E_*xy*_) were combined to calculate the RMSE of displacement and strain. Error bars are the standard error of the mean (n=7). Scale bar=10 mm.

Statistical results showed the NCS and CS groups were significantly different in both the raw displacement RMSE (compared to raw displacement average 8), smoothed strain RMSE (compared to smoothed strain average 8), and average. When comparing the raw displacement RMSE of the NCS average 5 data with the CS group, there was no significant difference observed in CS average 3 (Table 1). This indicates that CS average 3 has a similar RMSE with NCS average 5 which can save 40% of scanning time. For the smoothed strain RMSE, NCS average 4 was not significantly different with CS average 3 which takes 25% less time. Beyond average 6 and average 4 of raw displacement and smoothed strain of the NCS group, the CS group RMSE was significantly higher in all averages.

**Table 1.**
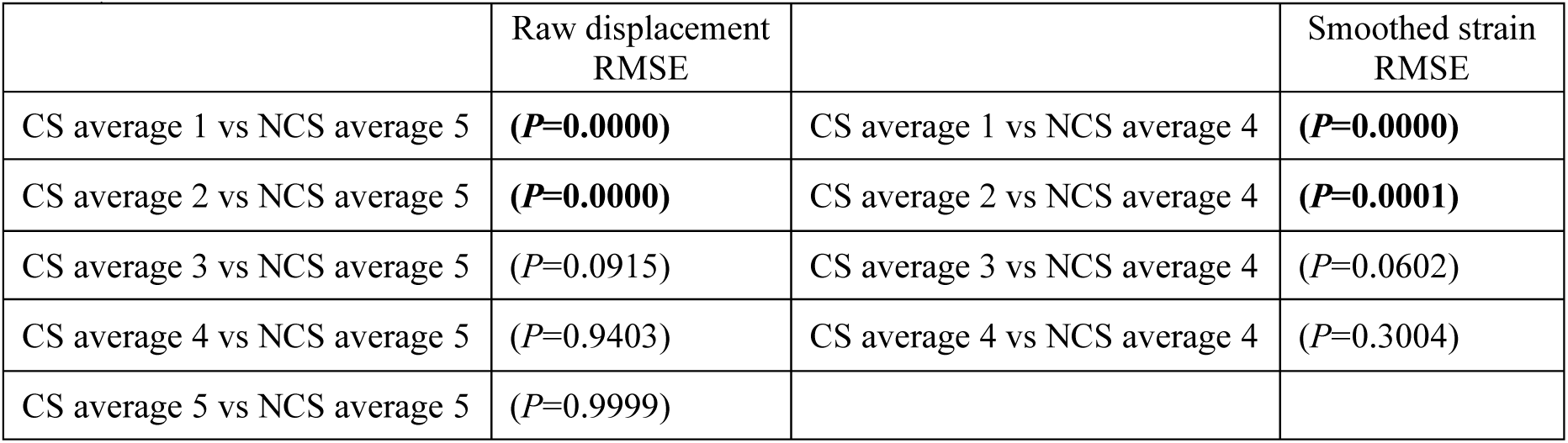
Comparison of NCS RMSE average 4, 5 with the CS RMSE average 1-5 groups. The CS average 1 and 2 were significantly different with the NCS average 5 group in raw displacement RSME while CS average 3-5 were not. For smoothed strain RMSE, CS average 1 and 2 groups were significantly different with the NCS average 4 group. *P* values were rounded to the fourth decimal and anything under 0.05 was considered significantly different. Statistical significance (*P*<0.05) is indicated in bold.

## DISCUSSION

To our knowledge, this is the first study to report results of pixel-level, high temporal resolution (40 ms) displacement and strain maps on a cohort of human cartilage obtained by spiral DENSE MRI and to demonstrate the potential of CS for accelerating DENSE MRI for displacement and strain quantification. We also measured cartilage T_1ρ_ and T_2_ values and examined changes in T_1ρ_ after varus loading. Spiral DENSE MRI successfully showed a gradual change of cartilage displacement and strain in response to varus load while CS yielded comparable results with images which could save utmost 25-40% of scanning time. The relaxometry values were comparable with studies conducted on healthy cartilage [39]–[41] while no significant change was observed in T_1ρ_ after repetitive varus loading, implying that the varus load was not detrimental to the knee joint [20]. Our analysis on cartilage validates the extraction of quantitative biomechanical measures in in vivo human subjects using spiral DENSE MRI and CS, demonstrating the potential of the routinal clinical usage and the development of imaging biomarkers for the detetion of early cartilage degeneration.

A previous study from our group [20] introduced using spiral DENSE MRI during compression on a phantom and ex vivo bovine joints while this paper expanded the methodology on in vivo humans, in essence showing the applicability of our method in clinical studies. The experimental settings we used in this study was adopted from the previous study since the dimensions of juvenile bovine joints and human knees were similar and the identical control system was used to operate the knee loading device. The implemented device for varus loading was completely MR safe and had precision and repeatability in terms of displacement smaller than the MR image pixel size, 360 µm [28]. The amount of varus load we selected, 0.5× BW on the medial cartilage, was in an acceptable range since the compressive load that human knee experiences while walking is 1.5–4× BW and 7–8× BW on extreme conditions like squatting [42], [43].

The acquired displacement and strain values on cartilage matched with visual observation during varus loading (Figure 2). Despite the translational motion occurring inside the knee coil, pronounced strain was detected in the strain maps. All images were captured in the coronal plane and the knee motion caused by varus load did not move out of the plane. The loading scheme applied (0.5 Hz) was adequate for physiologically relevant motion [44], [45]. Other approaches to measure cartilage strain in vivo caused by physical activities were to measure cartilage thickness before and after activities and studies have shown the maximum strain to be around 5% [46], [47]. This was less than what we found in this paper (9%). The difference in strain values could be that spiral DENSE MRI directly measured the displacement caused by the external load while cartilage thickness from other groups was measured after a physical activity was completed as a result, reflecting the creep deformation. Also, the ROI we selected on cartilage was capturing the contact area of femur and tibia together which differs with other studies where the cartilage thickness was measured on the tibia.

We found an interesting trend where there was significant difference of the spiral DENSE MRI results between male and female volunteers (Figure 3). One possible explanation for the greater displacement in *x* for male participants is the leg weight in females are higher regards to their total BW [48], [49]. Higher leg weight intuitively will require more load relative to BW to shift the knee and have the same displacement compared with male subjects. Also, male and females have a difference in femur and tibia axial alignment where females have a larger tibiofemoral angle forming more valgus knees than their male counterparts [50]–[52]. Therefore, the varus load we applied will act differently on female participant’s medial condyle, specifically which involves a knee alignment stage followed by pure varus load whereas male participants will experience less alignment stage and more varus load. Although there was no statistical difference found in strain between males and females, there was an interesting trend of strain decreasing during varus load which was only observed in females. This illustrates of how different bone structures can affect the loading cartilage receives during similar physical activities and implies why knee alignment is a risk factor for detecting OA [53], [54].

We measured T_1ρ_ before and after our cyclic loading protocol (Figure 4) since recent studies have reported a change of T_1ρ_ after force being applied to cartilage [20], [55], [56]. Some studies showed after mild physical activities, T_1ρ_ decreased due to the temporary loss of fluid [55], [56]. The varus test used in this study was solely compressing the medial condyle without any muscle activities and the temporary water content exudtion could have been minimum, not affecting the T_1ρ_ measurements. This aligned with what we found in our previous study where only in extreme conditions, T_1ρ_ started to change [20], and indicates that the varus load is a less harmful experiment to run. Consequently, this experiment could be adopted on knee OA patients who are limited in performing certain physical activties. T_2_ values were measured as a marker for cartilage water and collagen content, which is associated with tissue mechanical properties and reflects cartilage degradation [57]–[59]. Thus, the variation of strain measured by DENSE MRI could be linked to their T_2_ measurements. However, the T_2_ values measured were in a narrow range (SD 3.70 ms) and we found no correlation with the strain. Once this analysis is applied on a broad age group and participants with defected cartilage, a wider range of strain and T_2_ values will be anticipated where potential correlations can be found.

The effects of CS was substantial which greatly improved the image quality and considerably reduced the displacement and strain error acquired by spiral DENSE MRI (Figure 5-7). Shorter imaging time for our method led to fewer loading cycles being applied on the knee, which is advantageous when testing defected knees where excessive loading could be painful for the joint [60], [61]. Chen *et al* [62] has shown the potential of CS on spiral DENSE MRI of cardiac tissue where the measured displacement and circumferential strain error from under-sampled data improved after applying a CS method, BLOSM. In this paper, we assessed the CS effects using BART on different image averages which is readily controlled by the end-user. BART is an open-source online toolbox providing iterative reconstruction schemes on MR images [63]. The fact that we used BART with minimal fine-tunning and showed significant improvement in terms of image quality while spending much less imaging time on-line is promising. The reconstruction step in CS induced a denoising effect resulting in smooth and accurate displacement and strain fields which is coherent with other studies [64], [65]. However, CS contains an error caused by perturbation and the approximation in the model [66], [67] which reflected on the reconstructed images and accumulated over image averages. This is the cause of the high average CS data not improving the RMSE as much as the NCS data and not to asymptote at 0. Also, applying smoothing to CS displacement maps reduced the effects of CS which was later reflected on the smoothed strain RMSE plot where the NCS and CS group intersect while raw strain RMSE showed substantial improvements compared to NCS (Figure 7C). Hence, at low average and raw data the performance of CS was evident but the inherent CS error exacerbated over processing which reflected on the RMSE measurements.

In this study, there were some limitations that can be considered for future directions. The ROI we selected was on the tibiofemoral contact area combining the tibia and femur. To further analyze the biomechanics in the femur, tibia, and interface, separate ROIs should be used. Higher spatial resolution images would allow drawing separate ROIs and elucidate local spatial variations within the displacement maps which become less attenuated by smoothing. In addition, the CS algorithm can be further optimized to provide more accurate strain maps which show improvements across higher image averages. Lastly, comparing our method to kinematic analysis would provide a more robust understanding on cartilage degeneration related to physical activities. Motion capture methods [68]–[70] can quantitatively analyze human physical movements and is widely being used on in vivo studies. It would be meaningful to apply both analysis and find a correlation between the biomechanical responses of a joint with the kinematical information to observe how the analysis evolves on patients with different degeneration levels.

## CONCLUSION

In conclusion, this paper demonstrates using spiral DENSE MRI and CS on healthy human cartilage during varus loading. The obtained displacement and strain maps on the tibiofemoral contact region reveal the gradual shift over time. Also, there are significant differences between gender where male participants experience more displacement in the loading direction. The T_1ρ_ measurements before and after varus loading do not change significantly. Finally, the RMSE of displacement and strain after CS significantly decreases resulting in saving maximum 25-40% of scanning time. The findings in this study show spiral DENSE MRI with CS can be facilitated to clinical studies providing high spatial and temporal resolution mechanical responses of cartilage with reduced scanning times and less pain to participants. Future studies can be expanded to various age and OA grade groups for investigating relevant biomarkers that are sensitive to tissue degeneration.

## ACKNOWLEDGEMENTS

The authors would like to acknowledge funding from NIH R01 AR063712. S.E.S was funded through NIH T32 GM-065103. The authors are thankful to Teryn S. Wilkes for operating the MRI at Intermountain Neuroimaging Consortium located at University of Colorado, Boulder, and Nancy C. Emery for statistical consultation.

## AUTHOR CONTRIBUTIONS

Conceptualization, W.L., C.P.N.; Methodology & Data Collection, W.L., E.Y.M., H.Z.; Device Manufacturing, H.Z.; Statistical Analysis, W.L., S.E.S., D.A.R.; Writing – Original Draft, W.L., C.P.N., Writing – Review & Editing, All Authors; Funding Acquisition, C.P.N.

